# Body composition and growth in full-term small for gestational age and large for gestational age Swedish infants assessed with air displacement plethysmography at birth and at 3-4 months of age

**DOI:** 10.1101/468785

**Authors:** Anna Larsson, Peter Ottosson, Caroline Törnqvist, Elisabeth Olhager

## Abstract

**Background:** Being born small for gestational age (SGA) or large for gestational age (LGA) has short and long term metabolic consequences. There is a growing interest in the extent to which body composition, both in the short and the long term, differs in infants born at the extremes of these birth weights.

**Methods:** Body composition in 25 SGA and 25 LGA infants were assessed during the first days of life and at 3-4 months of age using air displacement plethysmography.

**Results:** SGA infants had significantly lower body fat (%) at birth compared to LGA infants. SGA infants increased their body weight and length at a significantly higher rate between birth and 3-4 months than LGA infants. Fat mass (g) in SGA infants increased 23 times between birth and 3-4 months of age compared to 2.8 times for LGA infants. At 3-4 months of age LGA infants reached a threshold in body fat (%) while SGA infants were still gaining body fat (%).

**Conclusion:** Several significant differences have been identified between SGA and LGA infants, indicating that the effects of intrauterine life continues to play an important role in body composition and growth during the first 3-4 months of life.

## Introduction

Unfavourable events during foetal life may lead to infants being born small for gestational age (SGA) or large for gestational age (LGA) which in turn is associated with adverse consequences for health later in life (1, 2, 3). Inadequate nutrient supply during foetal life caused by uteroplacental vascular insufficiency is the most common cause of being born SGA while being born LGA is associated with maternal diabetes mellitus, high maternal body mass index (BMI) and excessive weight gain during pregnancy (4, 5). The incidence of being born SGA and LGA varies in different parts of the world. According to Swedish statistics, in 2014 3.0 % of live-born infants weighed less than 2500 g and 3.7 % weighed more than 4500 g (6). The incidence of being LGA at birth has increased 15-20% during the last decades in several developed countries (7). Both SGA and LGA are associated with metabolic complications early and later in life (8, 9). Early in life there is a risk of low blood sugar for both LGA and SGA infants and later in life being born SGA is associated with an increased risk of developing adiposity and metabolic disease, particularly hypertension, increased cardiovascular mortality and type 2 diabetes mellitus (8, 9, 10). It is not clear whether these consequences are due to foetal programming caused by intrauterine malnutrition or the rapid weight gain early in life (8). Being born LGA is associated with type 1 diabetes later in life (10). Whether infants born LGA may be at risk of obesity and diabetes type 2 later in life is not clear.

There is a growing interest in the extent to which body composition, both in the short term and the long term, differs in infants and children born at the extremes of birth weights (3, 8). At birth, being LGA is associated with higher body weight and body fat (%) and slower postnatal weight gain compared to what is considered appropriate for gestational age (AGA) and being SGA is associated with lower body weight and body fat (%) at birth and higher postnatal weight gain compared to AGA (11, 12, 13).

Based on these findings, we hypothesised that there is a distinct difference between SGA and LGA infants in terms of body composition at birth and the subsequent growth in body weight, body length, fat mass, fat-free mass and body fat (%) during the first 3-4 months of life.

In recent years Pea Pod, based on air displacement plethysmography (ADP), has been launched as an accurate and reliable technique for measuring body composition in infants weighing up to 8 kg (18). Several studies in full-term neonates using the device have been published (16, 18).

We have carried out a longitudinal cohort study which aims to compare body composition and growth in full-term SGA and LGA infants during the first 3-4 months of life using ADP.

## Material and Methods

### Subjects

Two sample groups consisting of 25 SGA and 25 LGA infants respectively were recruited from the maternity ward at the University Hospital of Linköping, Sweden from January 2011 to April 2012. Inclusion criteria besides being SGA or LGA were birth at full term (gestational week 37-42 assessed using an ultrasound scan at approximately 12 weeks pregnancy), normal Apgar score, no malformations, no need for ventilation support or parenteral nutrition and completion of both first and second examinations.

At the first examination, 12 of the SGA infants were exclusively breast-fed and the remaining received both breast milk and formula. For LGA infants the corresponding figures were 13 and 12 respectively. At the second examination 13 of the SGA infants were exclusively breast-fed, 4 received both breast milk and formula, 7 received breast milk and solid foods, and 1 received formula and solid foods. The corresponding figures for LGA infants were 15,
8, 1, and 1 respectively.

### Study design

The study is based on two examinations: the first examination was performed during the first days of life before the mothers and infants were discharged from the maternity ward. The second examination was performed at 3-4 months of age. At birth the infants were weighed naked on an electronic baby scale (Tanita Corporation, Japan). Body length was measured on a length board and head circumference was measured with a non-elastic measuring band. At first and second examinations, ADP was used to measure body weight, fat mass, fat-free mass and body fat (%) and body length was measured on a length board and head circumference was measured with a non-elastic measuring band.

### Body composition

Body composition was assessed using Pea Pod Infant Body Composition System (LMI Concord, CA, USA) software 3.0.1. Its physical design and measurement procedures have been described elsewhere in detail (13). The Pea Pod has been validated and found to have high reliability and compliance to reference methods (15, 16). Using the principles of air displacement plethysmography, the Pea Pod system measures body volume, which together with body weight, is used to calculate body density. Body composition is then calculated using a two compartment model assuming a fat mass density of 0.9007 g/ml and using fat-free mass density values of Fomon et al. (17).

The measurement starts with the infant being weighed on an electronic scale that is part of the machine. Then the infant is placed without clothing in a supine position in the chamber which maintains a temperature of around 30° C. There is no need to avoid movement during the examination; hence the infant can be fully awake. The infant’s hair is kept flat using a tight cap provided by the supplier to avoid artifacts during measurement. The examination in the chamber takes two minutes and another two to four minutes before and after placing the infant in the chamber. All infants were examined by the same person (CT).

### Statistics

Data storage, calculations and analysis were performed using standard software IBM SPSS Statistics version 21, running on PC with Windows XP Professional SP3. Mean value and standard deviation were calculated using Compare Means. Linear Correlation was used to identify correlation coefficients and statistical significance. Stepwise Linear Regression was used to identify the most significant independent predicting factors. An Independent Samples t-test was used to verify statically significant differences between data-sets. Statistical significance was defined as p < 0.05.

### Ethical considerations

This study has been approved by the Human Research Ethics Committee of the Medical Faculty at Linköping University. All parents have given informed consent.

## Results

### First examination

Characteristics of the infants at birth are given in Table 1. SGA infants were born at 38.9 ±1.6 weeks while LGA infants were born at 40.1 ±1.5 weeks.

**Table 1.**
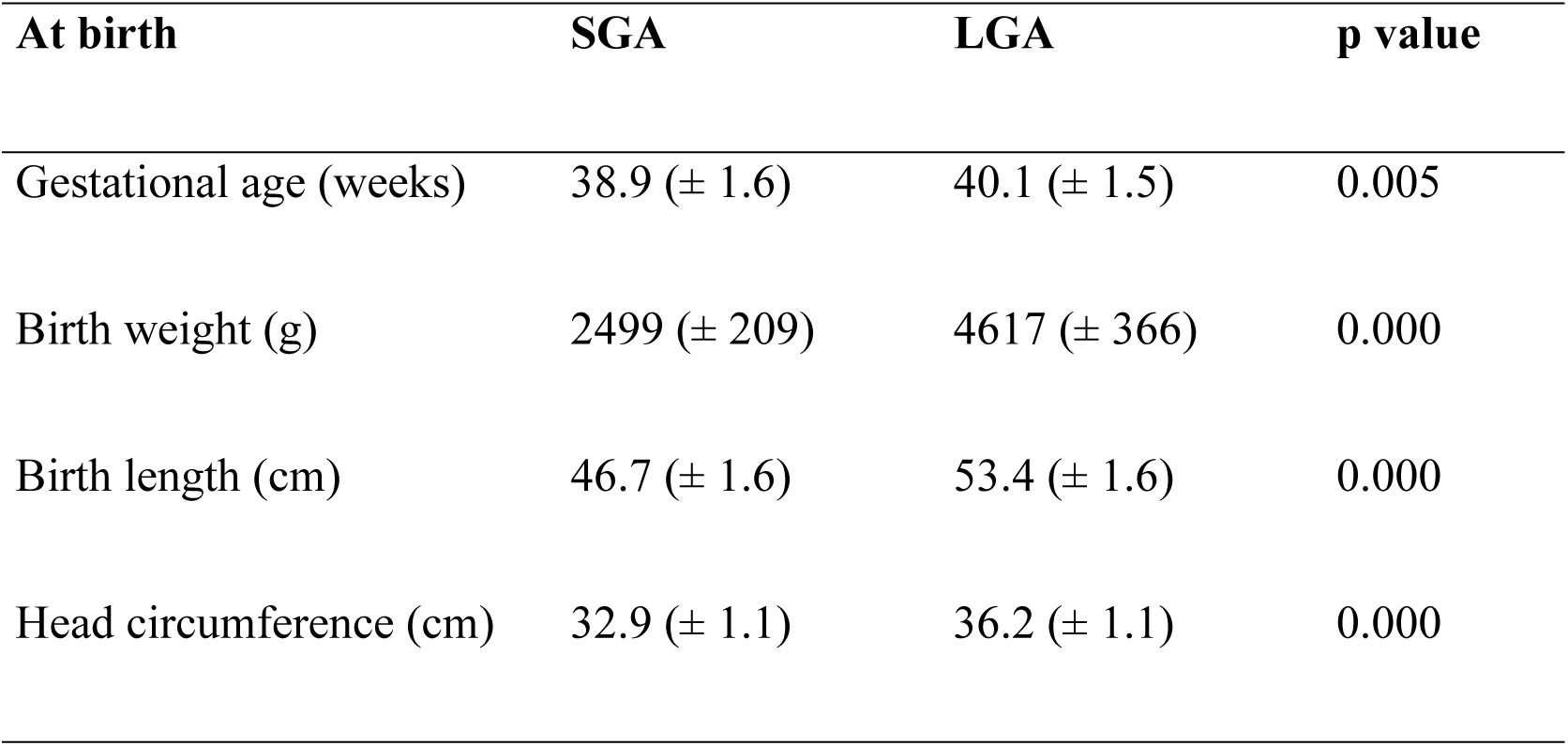
Gestational age, body weight, body length and head circumference of SGA infants (n = 25) and LGA infants (n = 25) at birth.

In Table 2 age, body length, body weight, body fat (%), fat mass and fat-free mass at first and second examinations are presented. SGA infants contained 3.7 ± 2.0 % body fat compared to LGA infants, who contained 17.3 ± 4.6 % body fat.

**Table 2.** Age, body length, body weight, body fat (%), fat mass and fat-free mass in SGA infants (n = 25) and LGA infants (n = 25) at the first and second examinations.

| **At 1<sup>st</sup> examination** | **SGA** | **LGA** | **p value** |
| --- | --- | --- | --- |
| Age (weeks) | 0.4 (± 0.3) | 0.4 (± 0.4) | NS |
| Length (cm) | 46.7 (± 1.6) | 53.4 (± 1.6) | 0.000 |
| Weight (g) | 2427 (± 225) | 4451 (± 414) | 0.000 |
| Body fat (%) | 3.7 (± 2.0) | 17.3 (± 4.6) | 0.000 |
| Fat mass (g) | 90 (± 46) | 773 (± 234) | 0.000 |
| Fat-free mass (g) | 2337 (± 233) | 3677 (± 361) | 0.000 |
| **At 2<sup>nd</sup> examination** | **SGA** | **LGA** | **p value** |
| Age (weeks) | 16.7 (± 2.1) | 14.0 (± 2.0) | 0.000 |
| Length (cm) | 60.7 (± 2.9) | 63.5 (± 1.9) | 0.000 |
| Weight (g) | 6031 (± 662) | 7162 (± 704) | 0.000 |
| Body fat (%) | 25.8 (± 4.4) | 27.6 (± 4.8) | NS |
| Fat mass (g) | 1574 (± 372) | 1978 (± 414) | 0.001 |
| Fat-free mass (g) | 4457 (± 413) | 5184 (± 586) | 0.000 |

NS = not significant

In LGA infants gestational age correlated with birth length (r = 0.755, p < 0.0001), birth weight (r = 0.757, p < 0.0001), fat-free mass (r = 0.481, p < 0.015) and fat mass (r = 0.492, p < 0.012).

### Second examination

At second examination SGA infants were significantly shorter than LGA infants and had significantly lower body weight. SGA infants also had significantly lower fat mass and fat-free mass. No significant difference was found in body fat (%) between the two groups. SGA infants contained 25.8 ± 4.4 % body fat and LGA infants contained 27.6 ± 4.8 % body fat.

### Growth

As shown in Table 3, SGA infants had significantly faster growth than LGA infants in body length, body weight, body fat (%), fat mass and fat-free mass between the first and second examinations. Figure 1 shows body weight, length, fat free and fat mass, body fat in % SGA and LGA infants at first and second examination.

**Table 3.** Growth and growth rate per week in body length, body weight, body fat (% units), fat mass and fat-free mass between the first and second examinations in SGA and LGA infants.

| **Growth** | **SGA** | **LGA** | **p value** |
| --- | --- | --- | --- |
| Length (cm) | 14.0 (± 3.0) | 10.1 (± 2.3) | 0.000 |
| Weight (g) | 3604 (± 633) | 2711 (± 645) | 0.000 |
| Body fat (% units) | 22.1 (± 4.8) | 10.3 (± 3.3) | 0.000 |
| Fat mass (g) | 1484 (± 372) | 1205 (± 356) | 0.009 |
| Fat-free mass (g) | 2120 (± 430) | 1507 (± 395) | 0.000 |
| **Growth rate** | **SGA** | **LGA** | **p value** |
| Length (cm/w) | 0.86 (± 0.14) | 0.75 (± 0.18) | 0.021 |
| Weight (g/w) | 223 (± 34) | 200 (± 39) | 0.028 |
| Body fat (% units/w) | 1.4 (± 0.3) | 0.8 (± 0.2) | 0.000 |
| Fat mass (g/w) | 92 (± 21) | 89 (± 23) | NS |
| Fat-free mass (g/w) | 132 (± 26) | 111 (± 26) | 0.006 |

NS = not significant

**Figure 1.**
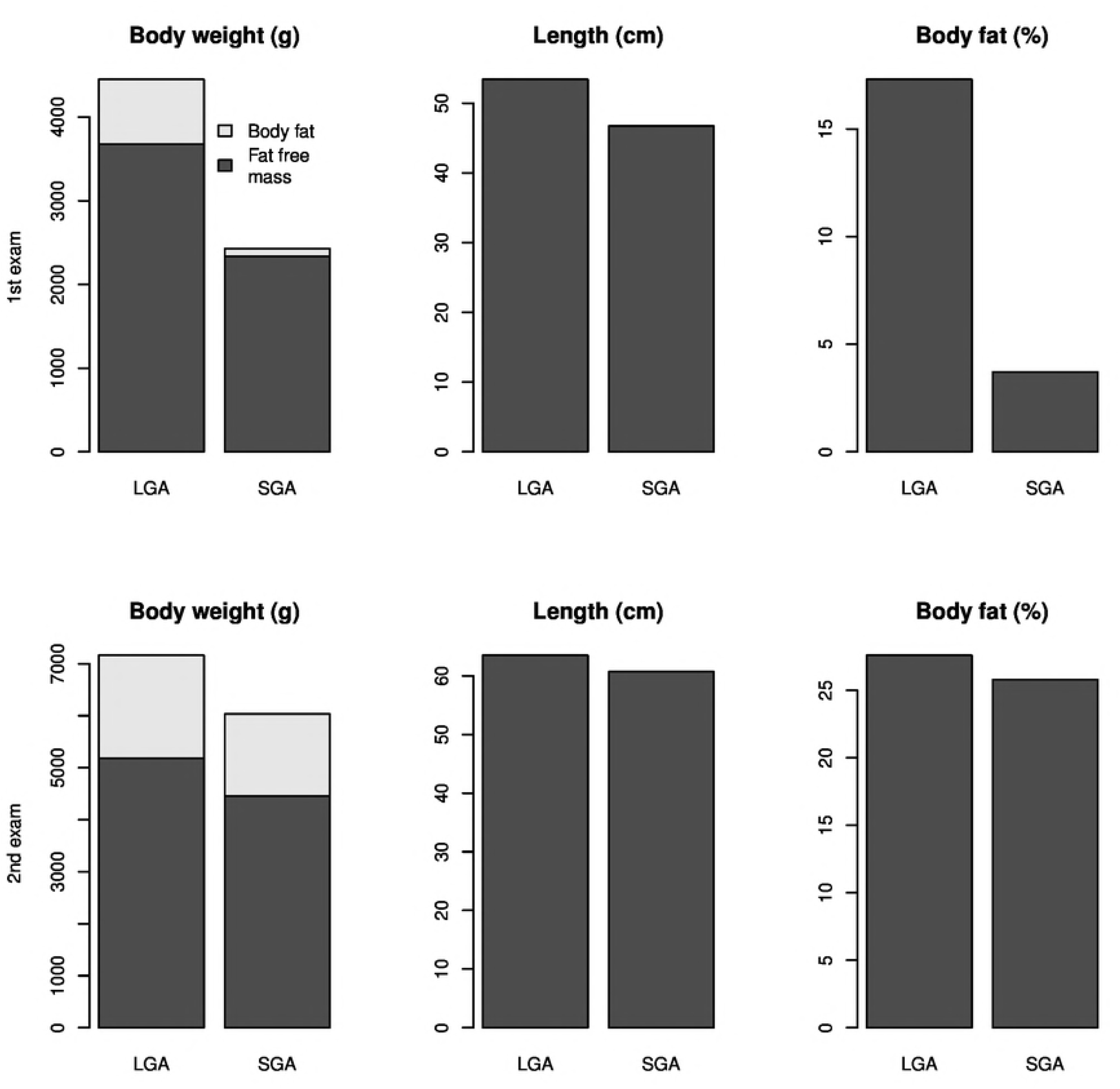
Body weight and length, fat free and fat mass and body fat in % in SGA and LGA infants at first and second examination

Since there was a significant difference in age at the second examination between SGA and LGA infants, LGA infants were measured at 14.0 (± 2.0) weeks while the corresponding figure for SGA infants was 16.7 (± 2.1). A linear model of growth rate in body weight, body length, body fat (%), fat mass and fat-free mass per week between birth and second examination was assumed, as shown in Table 3. There was a significant difference between the groups where SGA infants had a faster growth in body length, body weight, body fat (%) and fat-free mass per week than LGA infants. According to this model there was no significant difference in growth per week in fat mass between the groups. However, SGA infants increased their fat mass 23 times from first to second examination and LGA infants only 2.8 times (adjusted to the same mean age as SGA infants at second examination).

## Discussion

In this study we have explored the early changes in body composition in full term SGA and LGA infants using the ADP technique. Our main finding is that SGA infants have a significantly higher increase in body length, body weight, fat-free mass and body fat (%) compared with LGA infants between birth and 3-4 months of age. At first examination SGA infants were significantly shorter, lighter and had lower amounts of fat-free mass, fat mass and body fat (%) compared with LGA infants. At second examination SGA infants were still significantly shorter and lighter compared with LGA infants, and had significantly lower amount of fat mass and fat-free mass. However, there was no significant difference in body fat (%) between SGA and LGA infants and there was no significant difference between SGA and LGA infants in fat mass growth rate,

SGA infants increased their fat mass 23 times from first to second examination while the corresponding figure for LGA infants was 2.8 times (adjusted to the same age for second examination as SGA infants) reflecting the very low fat mass of SGA infants at birth compared to LGA infants. At birth LGA infants had on average 8.5 times the amount of fat mass compared to SGA infants in spite of the fact that the average body weight of LGA infants was only 1.8 times that of the SGA infants. The corresponding increase in fat mass in AGA infants is 4 times according to calculations based on data presented by Eriksson et al. (18). In this respect LGA infants behave more like AGA infants.

The growth rate in body weight, body length, fat-free mass and body fat (%) of SGA infants was high compared with LGA infants in this study. This has also been shown in other studies; for example Sun et al. state that fatter infants have a slower growth rate from birth to 5 months of age than thinner infants (11). They speculated on the mechanism behind the slower growth in LGA infants, hypothesising that both genetic susceptibility and exogenous factors such as breast feeding have an influence on the growth of fatter and heavier new-born infants (9). In another study, embedded in the Generation R study, Ay et al. showed that infants with catch-up growth in the first 6 weeks after birth had more body fat (%) at 6 months (19). Ong et al. speculate that during the early months of life, when feeding patterns are strongly influenced by the infant, and growth is regulated by nutrition, inherent patterns of increased or decreased appetite and satiety may return the infant towards its genetic growth trajectory (1). To some extent this could be an explanation of the slower weight gain among LGA infants in this study, since the majority of the LGA infants in this study were breast fed on demand. A majority of the SGA infants in this study were also breast fed on demand.

Wells et al. speculate that SGA infants, due to poor foetal growth, constrain lean body mass and thereby metabolic capacity, and the following rapid catch-up growth therefore leads to adiposity and increased metabolic load (20). In a study by Ibáñez et al. SGA infants show a characteristic catch-up growth compared to AGA infants, but remain shorter and develop a more central adiposity and higher degree of insulin resistance than AGA infants at age 2-4 years (9).

Both SGA and LGA infants keep growing in body length, body weight and fat-free mass until birth, shown as a positive correlation to gestational age at birth. SGA infants also keep growing in head circumference, while LGA infants seem to have reached a threshold, shown as no correlation between gestational age at birth and head circumference. Thomas et al. show that this threshold level in head circumference appears intrauterine at some time before 40 gestational weeks (21). The correlation between gestational age and head circumference of SGA infants in this study indicates that although full term, before birth the infants had been restrained from growing along their predetermined growth trajectory to reach a threshold in head circumference. In LGA infants gestational age at birth also shows a positive correlation to fat mass, which SGA infants do not, indicating that SGA infants suffer from inadequate nutrient supply. Eriksson et al. found no correlation between body fat (%) at birth and gestational age at birth for AGA infants (18), which is also true for SGA and LGA infants. In SGA infants, age in weeks is a major predicting factor for body fat (%) at second examination while in LGA infants, body fat (%) reached a threshold at the time of the second examination, shown as no correlation between age at second examination and body fat (%) for LGA infants. At 3-4 months of age SGA infants had caught up with LGA infants in terms of body fat (%). We speculate that intrauterine growth restriction initiates the driving force behind the dramatic relative growth in fat mass of SGA infants compared to LGA infants. This speculation is supported by the findings of a study by Ay et al. where it is shown that catch-down in weight in the third trimester was strongly associated with postnatal catch-up within six weeks of birth and a higher body fat (%) at six months of age (19). In LGA infants increase in body fat (%) is depressed at 3-4 months but in SGA it is not. The interesting question is, for how long and to what level does body fat (%) in SGA infants continue to rise? In a study by Roggero et al. preterm SGA and AGA infants showed no difference in body fat (%) at three months of corrected age and continued to increase in body fat (%) thereafter (23). In another study, Hediger et al. found that body fat (%) was higher for SGA than LGA infants at 2 to 47 months of age (24). This corresponds with the results of a study by Ong et al., where catch-up growth was predicted by factors relating to intrauterine growth restraint of foetal growth and where children who showed catch-up growth between zero and two years of age were fatter and had more central fat distribution at five years of age than other children (25). Our study showed an increased fat accretion in SGA infants compared with LGA infants. The mechanism behind the increased accretion of fat is not fully understood, but the findings are well described as the thrifty phenotype, leading to increased risk for a number of conditions, such as impaired glucose tolerance, insulin resistance and relative adiposity, factors that are associated with coronary vascular disease, hypertensive diseases and diabetes as adults in the long term (26, 27). A theoretical possibility is that intrauterine growth restrictions that lead to SGA also lead to altered epigenetic programming (altered DNA methylation) of the infant. Tobi et al., however, did not find any significant difference in DNA methylation between preterm SGA and AGA infants in the specific loci they examined (28).

In conclusion, SGA infants grow faster in body weight, body length and fat-free mass compared to LGA infants between birth and 3-4 months of age. SGA infants also show a conspicuous stable growth in fat mass. In SGA infants we also found a positive correlation between the rate of increase in body fat (%) and gestational age at birth, as well as a clear pattern of sex-related catch-up growth, where males showed a stronger growth in fat-free mass than females.

## Acknowledgements

We wish to express our gratitude to the infants who participated in this study and to their parents.

## Authors’ contribution

AL and PO analysed data and prepared the manuscript. CT examined all the infants. EO planed the study, analysed the data and prepared the manuscript. At the time of the examination EO was head of the neonatal intensive care unit in Linköping

## Abbreviations

SGA: small for gestational age
LGA: Large for gestational age
ADP: air displacement plethysmography

